## Supplementary Material for "Deciphering the regulatory genome of *Escherichia coli*, one hundred promoters at a time"

### Supplementary Information for "Deciphering the regulatory genome of *Escherichia coli*, one hundred promoters at a time"

William T. Ireland[1], Suzannah M. Beeler[2] Emanuel Flores-Bautista[2], Nathan M. Belliveau[2,†], Michael J. Sweredoski[3], Annie Moradian[3], Justin B. Kinney[4], Rob Phillips[1,2,5,*]

[1] Department of Physics, California Institute of Technology, Pasadena, CA 91125
[2] Division of Biology and Biological Engineering, California Institute of Technology, Pasadena, CA 91125
[3] Proteome Exploration Laboratory, Beckman Institute, California Institute of Technology, Pasadena, CA 91125
[4] Simons Center for Quantitative Biology, Cold Spring Harbor Laboratory, Cold Spring Harbor, NY 11724
[5] Department of Applied Physics, California Institute of Technology, Pasadena, CA 91125
[†] Present address: Howard Hughes Medical Institute and Department of Biology, University of Washington, Seattle, WA 98195

### 1 Extended details of experimental design

#### 1.1 Choosing target genes

Genes in this study were chosen to cover several different categories. 29 genes had some information on their regulation already known to validate our method under a number of conditions. 37 were chosen because the work of [1] demonstrated that gene expression changed significantly under different growth conditions. A handful of genes such as *minC*, *maoP*, or *fdhE* were chosen because we found either their physiological significance interesting, as in the case of the cell division gene *minC* or that we found the gene regulatory question interesting, such for the intra-operon regulation demonstrated by *fdhE*. The remainder of the genes were chosen because they had no regulatory information, often had minimal information about the function of the gene, and had an annotated transcription start site (TSS) in RegulonDB.

#### 1.2 Choosing transcription start sites

A known limitation of the experiment is that the mutational window is limited to 160 bp. As such, it is important to correctly target the mutation window to the location around the most active TSS. To do this we first prioritized those TSS which have been extensively experimentally validated and catalogued in RegulonDB. Secondly we selected those sites which had evidence of active transcription from RACE experiments [2] and were listed in RegulonDB. If the intergenic region was small enough, we covered the entire region with our mutation window. If none of these options were available, we used computationally predicted start sites.

#### 1.3 Sequencing

All sequencing was carried out by either the Millard and Muriel Jacobs Genetics and Genomics Laboratory at Caltech (HiSeq 2500) on a 100 bp single read flow cell or using the sequencing services from NGX Bio on a 250 bp or 150 base paired end flow cell. The total library was first sequenced by PCR amplifying the region containing the variant promoters as well as the corresponding barcodes. This allowed us to uniquely associate each random 20 bp barcode with a promoter variant. Any barcode which was associated with a promoter variant with insertions or deletions was removed from further analysis. Similarly, any barcode that was associated with multiple promoter variants was also removed from the analysis. The paired end reads from this sequencing step were then assembled using the FLASH tool [3]. Any sequence with PHRED score less than 20 was removed using the FastX toolkit. Additionally, when sequencing the initial library, sequences which only appear in the dataset once were not included in further analysis in order to remove possible sequencing errors.

For all the MPRA experiments, only the region containing the random 20 bp barcode was sequenced, since the barcode can be matched to a specific promoter variant using the initial library sequencing run described above. For a given growth condition, each promoter yielded 50,000 to 500,000 usable sequencing reads. Under some growth conditions, genes were not analyzed further if they did not have at least 50,000 reads.

To determine which base pair regions were statistically significant a 99% confidence interval was constructed using the MCMC inference to determine the uncertainty.

#### 1.4 Growth conditions

The growth conditions studied in this study were inspired by [1] and include differing carbon sources such as growth in M9 with 0.5% Glucose, M9 with acetate (0.5%), M9 with arabinose (0.5%), M9 with Xylose (0.5%) and arabinose (0.5%), M9 with succinate (0.5%), M9 with fumarate (0.5%), M9 with Trehalose (0.5%), and LB. In each case cell harvesting was done at an OD of 0.3. These growth conditions were chosen so as to span a wide range of growth rates, as well as to illuminate any carbon source specific regulators.

We also used several stress conditions such as heat shock, where cells were grown in M9 and were subjected to a heat shock of 42 degrees for 5 minutes before harvesting RNA. We grew in low oxygen conditions. Cells were grown in LB in a container with minimal oxygen, although some will be present as no anaerobic chamber was used. This level of oxygen stress was still sufficient to activate FNR binding, and so activated the anaerobic metabolism. We also grew cells in M9 with Glucose and 5mM sodium salycilate.

Growth with zinc was preformed at a concentration of 5mM $ZnCl_2$ and growth with iron was preformed by first growing cells to an OD of 0.3 and then adding $FeCL_2$ to a concentration of 5mM and harvesting RNA after 10 minutes. Growth without cAMP was accomplished by the use of the JK10 strain which does not maintain its cAMP levels.

All knockout experiment were preformed in M9 with Glucose except for the knockouts for *arcA*, *hdfR*, and *phoP* which were grown in LB.

### 2 Validating Reg-Seq against previous methods and results

The work presented here is effectively a third-generation of the use of Sort-Seq methods for the discovery of regulatory architecture. The primary difference between the present work and previous generations [4, 5] is the use of RNA-Seq rather than fluorescence and cell sorting as a readout of the level of expression of our promoter libraries. As such, there are many important questions to be asked about the comparison between the earlier methods and this work. We attack that question in several ways. First, as shown in Figure S1, we have performed a head-to-head comparison of the two approaches to be described further in this section. Second, as shown in the next section, our list of candidate promoters included roughly 20% for which the community has some knowledge of their regulatory architecture. In these cases, we examined the extent to which our methods recover the known features of regulatory control about those promoters.

#### 2.1 Comparison between Reg-Seq by RNA-Seq and fluorescent sorting

As the basis for comparing the results of the fluorescence-based Sort-Seq approach with our RNA-Seq-based approach, we use information footprints, expression shifts and sequence logos as our metrics. Figure S1 shows examples of this comparison for four distinct genes of interest. Figure S1(A) shows the results of the two methods for the *lacZYA* promoter with special reference to the CRP binding site. Both the information footprint and the sequence logo identify the same binding site.

Figure S1(B) provides a similar analysis for the *dgoRKADT* promoter where once again the information footprints and the sequence logos from the two methods are in reasonable accord. Figure S1(C) provides a quantitative dissection of the *relBE* promoter which is repressed by RelBE. Here we use both information footprints and expression shifts as a way to quantify the significance of mutations to different binding sites across the promoter. Finally, Figure S1(D) shows a comparison of the two methods for the *marRAB* promoter. The two approaches both identify a MarR binding site.

#### 2.2 Ability of Reg-Seq to recover known regulatory architectures

In total, we have tested over 20 genes for which there is already some substantial regulatory knowledge reported in the literature. The successes and failures of this test are detailed in Figure S2. For those promoters which have strong evidence of a binding site, as determined by RegulonDB [6], we recover all relevant transcription factor binding sites for 12 out of 16 cases, the majority of relevant binding sites for 2 out of 16 cases, and miss all or most of the regulation for just 2 promoters. We identify a total of 22 previously known high evidence binding sites.

These results showcase that our method largely agrees with the established literature but also highlights several areas in which our method is prone to missing regulatory elements. One failure mode is caused by the presence of strong secondary binding sites. For example, in the *araC* promoter, as shown in Figure S2(C), the only binding signatures that appear in the information footprint are from a secondary RNAP site. The secondary site seems to be expressed constitutively, and in the cases where the primary start site is even partially repressed, the secondary start site will dominate transcription and obscure the many binding sites that are in this promoter.

If there are large numbers of regulatory elements, the data will often only show the few most important elements. If we look at the *marR* promoter in Figure S2(C), we can only see the signature of the two MarR sites even though CpxR, Fis, and CRP are all known to bind to the promoter. MarR is a strong enough repressor that mutating any of the other transcription factor sites is unlikely to meaningfully change gene expression unless the MarR site is also mutated. This illustrates that the regulatory architectures discovered in this study represent a lower bound on what exists in each promoter.

Finally, for some genes such as *dicA* there was no known TSS prior to the experiment. Although there is a small regulatory region between *dicA* and its neighboring gene, this does not ensure that we will include the strongest RNAP sites. Better mapping of transcription start sites could improve our method.

We next consider low evidence binding sites. Other research determined the locations of the low evidence sites through gene expression analysis and sequence comparison to consensus sequences [7, 8, 9]. For 5 promoters in our list, the binding sites location itself is not known, only that the TF in question regulates the gene. For these promoters we recover the known regulation in only 2 out of 15 cases. Comparison to consensus sequences can be unreliable and generate false positives when the entirety of the *E. coli* genome is considered. Gene expression analysis

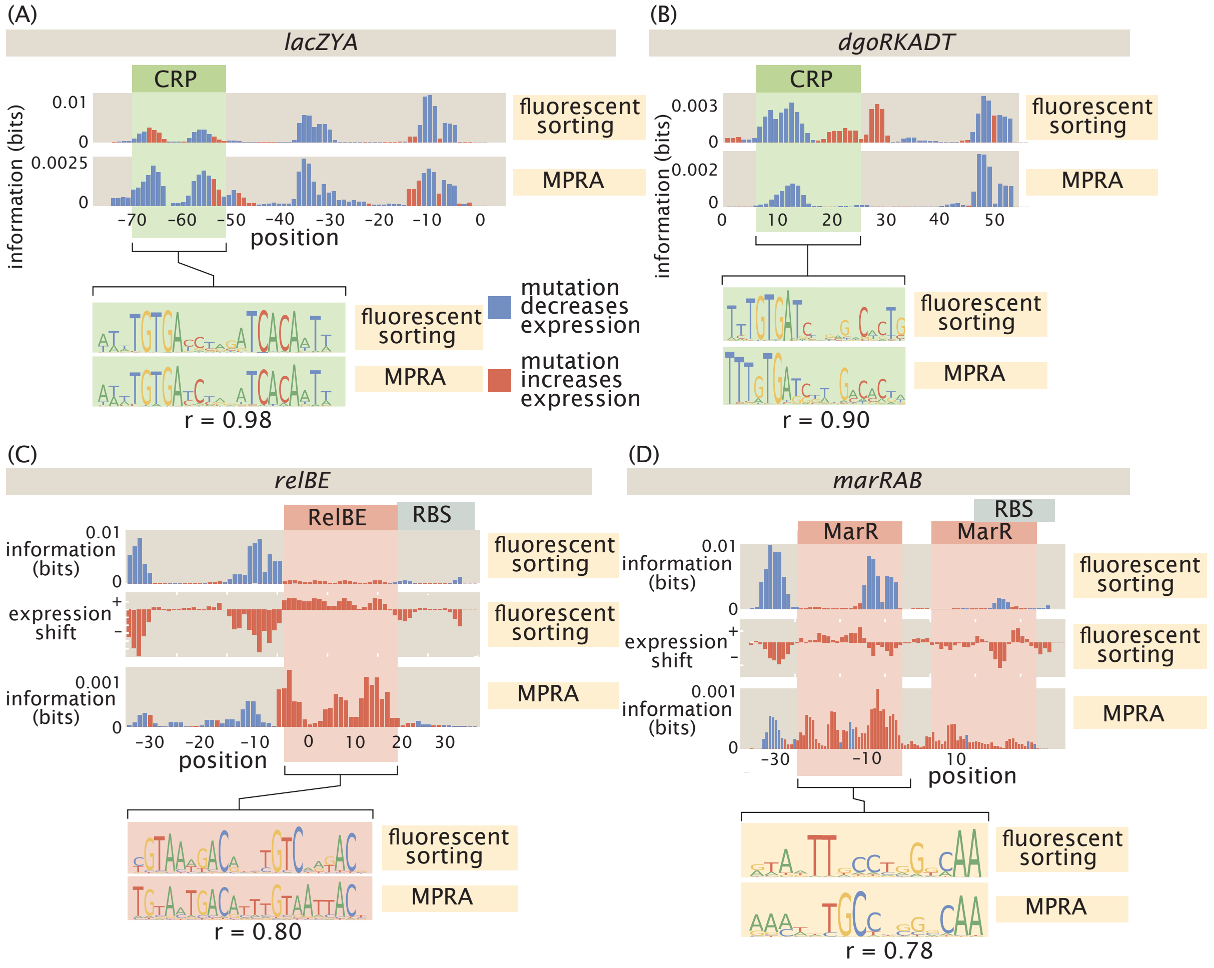

Figure S1: A summary of four direct comparisons of measurements using fluorescence and sorting and using RNA-Seq. (A) CRP binds upstream of RNAP in the *lacZYA* promoter. Despite the different measurement techniques for the two inferred energy matrices and their corresponding sequence logos, the CRP binding sites have a Pearson correlation coefficient of $r = 0.98$. (B) The *dgoRKADT* promoter is activated by CRP in the presence of galactonate. The FACS measurements were taken in the JK10 strain in the presence of 500mM cAMP. In both cases, a type II activator binding site can be identified based on the signals in the information footprint in the area indicated in green. Additionally the quantitative agreement between the CRP binding preference matrices are strong, with $r = 0.9$. (C) The *relBE* promoter is repressed by RelBE. The inferred matrices between the two measurement methods have $r = 0.8$. (D) The *marRAB* promoter is repressed by MarR. The features we can observe in the information footprint reflect this under measurement with both FACS or RNAseq. The inferred energy matrices (data not shown) and sequence logos shown have $r = 0.78$. The right most MarR site overlaps with a ribosome binding site. The overlap has a stronger obscuring effect on the sequence specificity of the FACS measurement, which measures protein levels directly, than it does on the output of the RNAseq measurement.

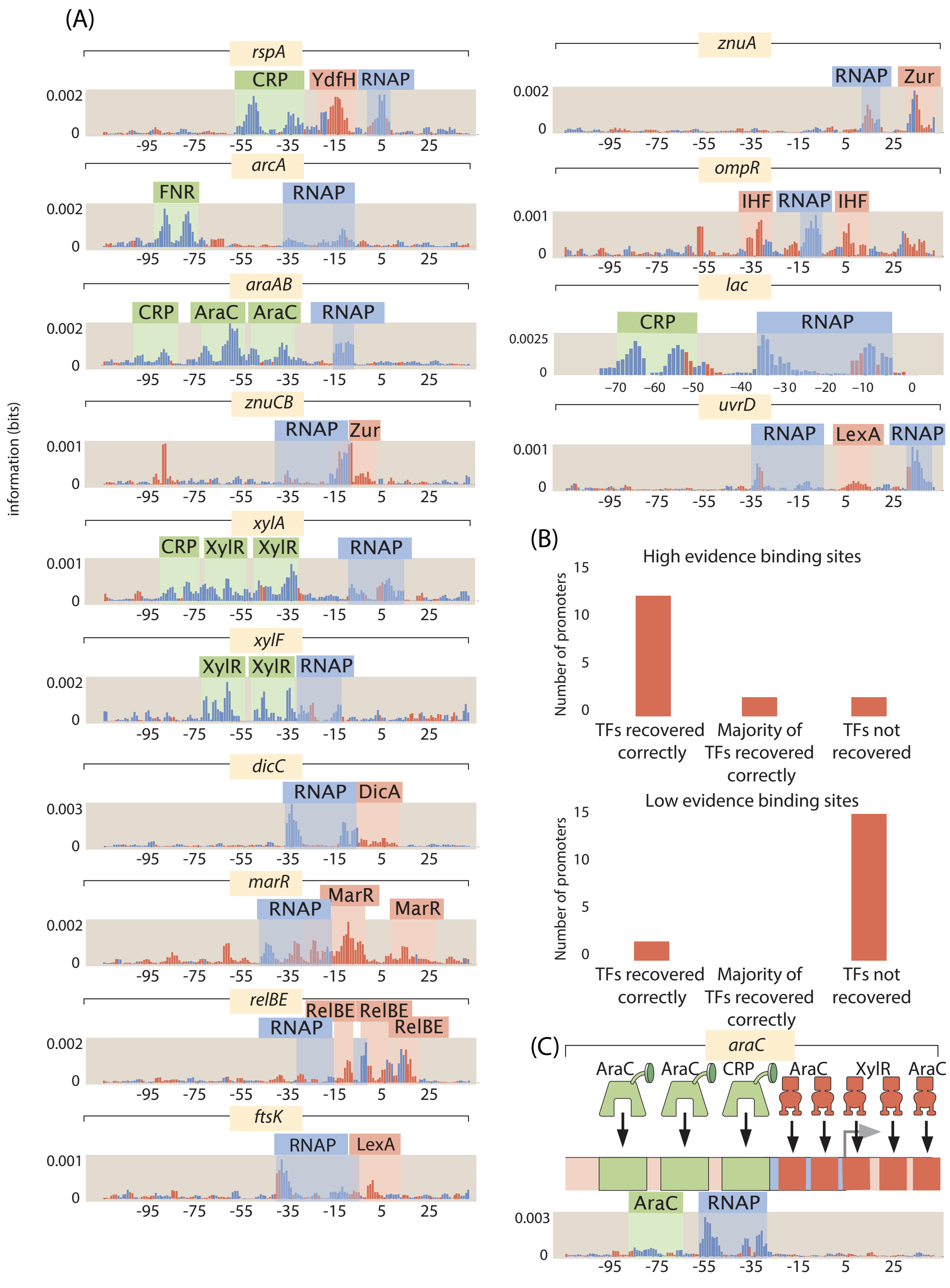

Figure S2: Reg-Seq analysis of "gold standard" promoters. (A) Information footprints for known and properly recovered binding sites. (B) A summary of how well the Reg-Seq results conform to literature results. The sites that are low evidence in the literature are determined by RegulonDB [6]. (C) The information footprint and known binding sites for the *araC* promoter. Despite all the binding sites present, the only binding signature that appears is for RNAP.

alone has difficulty ruling out indirect effects of a given transcription factor on gene expression and regulation determined by this method may occur outside of the 160 bp mutation window we consider. As our results recover high evidence sites well, the poor recovery of sites based on sequence gazing and gene expression analysis most likely indicates that these methods are unreliable for determining binding locations.

We note that the first aim of our methods is regulatory discovery. We would like to be able to determine how previously uncharacterized promoters are regulated and ultimately, this is a question of binding-site and transcription factor identification. For that task, we do not require perfect correspondence between the two methods. With regulatory sites identified, our next objective is the determination of energy matrices that will allow us to turn binding site strength into a tunable knob that can nearly continuously tune the strength of transcription factor binding, thus altering gene expression in predictable ways as already shown in our earlier work [10]. The r-values between energy matrices range from 0.78 to 0.96, indicating reasonable to very good agreement. Reg-Seq appears to be, if anything, more accurate than previous methods as it has higher relative information content in known areas of transcription factor binding and also does not have repressor-like bases on CRP sites as in Figure S1(A) and (B).

### 3 Extended details of analysis methods

#### 3.1 Information footprints

We use information footprints as a tool for hypothesis generation to identify regions which may contain transcription factor binding sites. In general, a mutation within a transcription factor site is likely to severely weaken that site. We look for groups of positions where mutation away from wild type has a large effect on gene expression. Our data sets consist of nucleotide sequences, the number of times we sequenced the construct in the plasmid library, and the number of times we sequenced its corresponding mRNA. A simplified data set on a 4 nucleotide sequence then might look like

| Sequence | Library Sequencing Counts | mRNA Counts |
|---|---|---|
| ACTA | 5 | 23 |
| ATTA | 5 | 3 |
| CCTG | 11 | 11 |
| TAGA | 12 | 3 |
| GTGC | 2 | 0 |
| CACA | 8 | 7 |
| AGGC | 7 | 3 |

One possible calculation to measure the impact of a given mutation on expression is to take all sequences which have base $b$ at position $i$ and determine the number of mRNAs produced per read in the sequencing library. By comparing the values for different bases we could determine how large of an effect mutation has on gene expression. However, in this paper we will use mutual information to quantify the effect of mutation, as [4] demonstrated could be done successfully. In Table 1 the frequency of the different nucleotides in the library at position 2 is 40% A, 32% C,

14% G and 14% T. Cytosine is enriched in the mRNA transcripts over the original library, as it now composes 68% of all mRNA sequencing reads while A, G, and T only compose only 20%, 6%, and 6% respectively. Large enrichment of some bases over others occurs when base identity is important for gene expression. We can quantify how important using the mutual information between base identity and gene expression level. Mutual information is given at position $i$ by

$$I_b = \sum_{m=0}^{1} \sum_{\mu=0}^{1} p(m,\mu) \log_2 \left( \frac{p(m,\mu)}{p_{mut}(m) p_{expr}(\mu)} \right). \tag{1}$$

$p_{mut}(m)$ in equation 1 refers to the probability that a given sequencing read will be from a mutated base. $p_{expr}(\mu)$ is a normalizing factor that gives the ratio of the number of DNA or mRNA sequencing counts to total number of counts.

The mutual information quantifies how much a piece of knowledge reduces the entropy of a distribution. At a position where base identity matters little for expression level, there would be little difference in the frequency distributions for the library and mRNA transcripts. The entropy of the distribution would decrease only by a small amount when considering the two types of sequencing reads separately.

We are interested in quantifying the degree to which mutation away from a wild type sequence affects expression. Although their are obviously 4 possible nucleotides, we can classify each base as either wild-type or mutated so that $b$ in equation 1 represents only these two possibilities.

If mutations at each position are not fully independent, then the information value calculated in equation 1 will also encode the effect of mutation at correlated positions. If having a mutation at position 1 is highly favorable for gene expression and is also correlated with having a mutation at position 2, mutations at position 2 will also be enriched amongst the mRNA transcripts. Position 2 will appear to have high mutual information even if it has minimal effect on gene expression. Due to the DNA synthesis process used in library construction, mutation in one position can make mutation at other positions more likely by up to 10 percent. This is enough to cloud the signature of most transcription factors in an information footprint calculated using equation 1.

We need to determine values for $p_i(m|\mu)$ when mutations are independent, and to do this we need to fit these quantities from our data. We assert that

$$\langle mRNA \rangle \propto e^{-\beta E_{eff}} \tag{2}$$

is a reasonable approximation to make. $\langle mRNA \rangle$ is the average number of mRNAs produced by that sequence for every cell containing the construct and $E_{eff}$ is an effective energy for the sequence that can be determined by summing contributions from each position in the sequence. There are many possible underlying regulatory architectures, but to demonstrate that our approach is reasonable let us first consider the simple case where there is only a RNAP site in the studied region. We can write down an expression for average gene expression per cell as

$$\langle mRNA \rangle \propto p_{bound} \propto \frac{\frac{p}{N_{NS}} e^{-\beta E_P}}{1 + \frac{p}{N_{NS}} e^{-\beta E_P}} \tag{3}$$

Where $p_{bound}$ is the probability that the RNAP is bound to DNA and is known to be proportional to gene expression in *E. coli* [11], $E_P$ is the energy of RNAP binding, $N_{NS}$ is the number of nonspecific DNA binding sites, and $p$ is the number of RNAP. If RNAP binds weakly then $\frac{p}{N_{NS}}e^{-\beta E_P} << 1$. We can simplify equation 3 to

$$\langle mRNA \rangle \propto e^{-\beta E_P}. \tag{4}$$

If we assume that the energy of RNAP binding will be a sum of contributions from each of the positions within its binding site then we can calculate the difference in gene expression between having a mutated base at position $i$ and having a wild type base as

$$\frac{\langle mRNA_{WT_i} \rangle}{\langle mRNA_{Mut_i} \rangle} = \frac{e^{-\beta E_{P_{WT_i}}}}{e^{-\beta E_{P_{Mut_i}}}} \tag{5}$$

$$\frac{\langle mRNA_{WT_i} \rangle}{\langle mRNA_{Mut_i} \rangle} = e^{-\beta (E_{P_{WT_i}} - E_{P_{Mut_i}})}. \tag{6}$$

In this example we are only considering single mutation in the sequence so we can further simplify the equation to

$$\frac{\langle mRNA_{WT_i} \rangle}{\langle mRNA_{Mut_i} \rangle} = e^{-\beta \Delta E_{P_i}}. \tag{7}$$

We can now calculate the base probabilities in the expressed sequences. If the probability of finding a wild type base at position $i$ in the DNA library is $p_i(m = WT|\mu = 0)$ then

$$p_i(m = WT|\mu = 1) = \frac{p_i(m = WT|\mu = 0)\frac{\langle mRNA_{WT_i} \rangle}{\langle mRNA_{Mut_i} \rangle}}{p_i(m = Mut|\mu = 0) + p_i(m = WT|\mu = 0)\frac{\langle mRNA_{WT_i} \rangle}{\langle mRNA_{Mut} \rangle}} \tag{8}$$

$$p_i(m = WT|\mu = 1) = \frac{p_i(m = WT|\mu = 0)e^{-\beta \Delta E_{P_i}}}{p_i(m = Mut|\mu = 0) + p_i(m = WT|\mu = 0)e^{-\beta \Delta E_{P_i}}}. \tag{9}$$

Under certain conditions, we can also infer a value for $p_i(m|\mu = 1)$ using a linear model when there are any number of activator or repressor binding sites. We will demonstrate this in the case of a single activator and a single repressor, although a similar analysis can be done when there are greater numbers of transcription factors. We will define $P = \frac{p}{N_{NS}}e^{-\beta E_P}$. We will also define $A = \frac{a}{N_{NS}}e^{-\beta E_A}$ where $a$ is the number of activators, and $E_A$ is the binding energy of the activator. We will finally define $R = \frac{r}{N_{NS}}e^{-\beta E_R}$ where $r$ is the number of repressors and $E_R$ is the binding energy of the repressor. We can write

$$\langle mRNA \rangle \propto p_{bound} \propto \frac{P + PAe^{-\beta \epsilon_{AP}}}{1 + A + P + R + PAe^{-\beta \epsilon_{AP}}} \tag{10}$$

If activators and RNAP bind weakly but interact strongly, and repressors bind very strongly, then we can simplify equation 10. In this case $A << 1$, $P << 1$, $PAe^{-\epsilon_{AP}} >> P$, and $R >> 1$. We can then rewrite equation 10 as

$$\langle mRNA \rangle \propto \frac{PAe^{-\beta\epsilon_{AP}}}{R} \tag{11}$$
$$\langle mRNA \rangle \propto e^{-\beta(-E_P - E_A + E_R)} \tag{12}$$

As we typically assume that RNAP binding energy, activator binding energy, and repressor binding can all be represented as sums of contributions from their constituent bases, the combination of the energies can be written as a total effective energy $E_{eff}$ which is a sum of contributions from all positions within the binding sites.

We fit the parameters for each base using a Markov Chain Monte Carlo Method. Two MCMC runs are conducted using randomly generated initial conditions. We require both chains to reach the same distribution to prove the convergence of the chains. We do not wish for mutation rate to affect the information values so we set the $p(WT) = p(Mut) = 0.5$ in the information calculation. The information values are smoothed by averaging with neighboring values.

#### 3.2 Analysis of mass spectrometry results

Mass spectrometry results were processed using MaxQuant [12] [13]. Spectra were searched against the UniProt *E. coli* K-12 database as well as a contaminant database (256 sequences). LysC was specified as the digestion enzyme. Proteins were considered if they were known to be transcription factors, or were predicted to bind DNA (using gene ontology term GO:0003677, for DNA-binding in BioCyc).

#### 3.3 Uncertainty due to number of independent sequences

1400 promoter variants were ordered from TWIST Bioscience for each promoter studied. Due to errors in synthesis, additional mutations are introduced into the ordered oligos. As a result, the final number of variants received was an average of 2200 per promoter. To test whether the number of promoter variants is a significant source of uncertainty in the experiment we computationally reduced the number of promoter variants used in the analysis of the *zapAB* -10 RNAP region. Each sub-sampling was performed 3 times. The results, as displayed in Figure S3, show that there is only a small effect on the resulting sequence logo until the library has been reduced to approximately 500 promoter variants.

#### 3.4 TOMTOM motif comparison

In some cases, we used an alternative approach to mass spectrometry to discover the TF identity regulating a given promoter based on sequence analysis using a motif comparison tool. TOMTOM [14] is a tool that uses a statistical method to infer if a putative motif resembles any previously discovered motif in a database. Of interest, it accounts for all possible offsets between the motifs. Moreover, it uses a suite of metrics to compare between motifs such as Kullback-Leibler divergence, Pearson correlation, euclidean distance, among others.

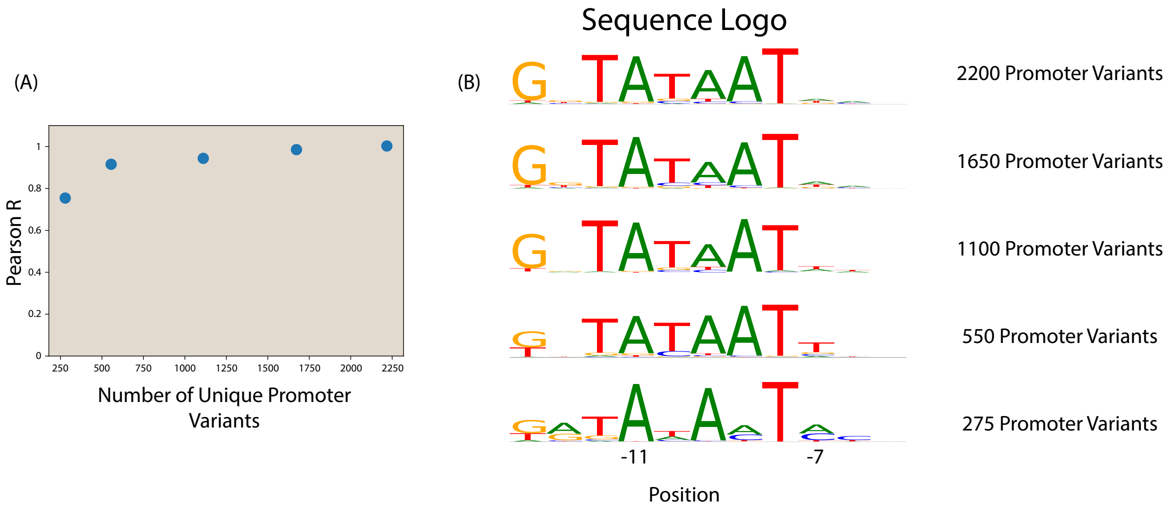

Figure S3: A comparison of RNAP -10 site sequence logos. (A) This figure shows the Pearson correlation coefficient between the energy matrix models inferred from the full dataset (2200 unique promoter variants) and that from a computationally restricted dataset. (B) Sequence logos of the RNAP -10 region from each sub-sampled dataset.

We performed comparisons of the motifs generated from our energy matrices to those generated from all known transcription factor binding sites in RegulonDB. Figure S4 shows a result of TOMTOM, where we compared the motif derived from the -35 region of the *ybjX* promoter and found a good match with the motif of PhoP from RegulonDB.

The information derived from this approach was then used to guide some of the TF knockout experiments, in order to validate its interaction with a target promoter characterized by the loss of the information footprint. Furthermore, we also used TOMTOM to search for similarities between our own database of motifs, in order to generate regulatory hypotheses in tandem. This was particularly useful when looking at the group of GlpR binding sites found in this experiment.

### 4 Additional results

#### 4.1 Binding sites regulating divergent operons

In addition to discovering new binding sites, we have discovered additional functions of known binding sites. In particular, in the case of *bdcR*, the repressor for the divergently transcribed gene *bdcA* [15], is also shown to repress *bdcR* in Figure S5(A). Similarly in Figure S5(B) IvlY is shown to repress *ilvC* in the absence of inducer. Divergently transcribed operons that share regulatory regions are plentiful in *E. coli*, and although there are already many known examples of transcription factor binding sites regulating several different operons, there are almost certainly many examples of this type of transcription that have yet to be discovered.

Multi-purpose binding sites allow for more genes to be regulated with fewer binding sites. However, they can also serve to sharpen the promoter's response to environmental cues. In the case of *ilvC*, IlvY is known to activate *ilvC* in the presence of inducer. However, we now see that

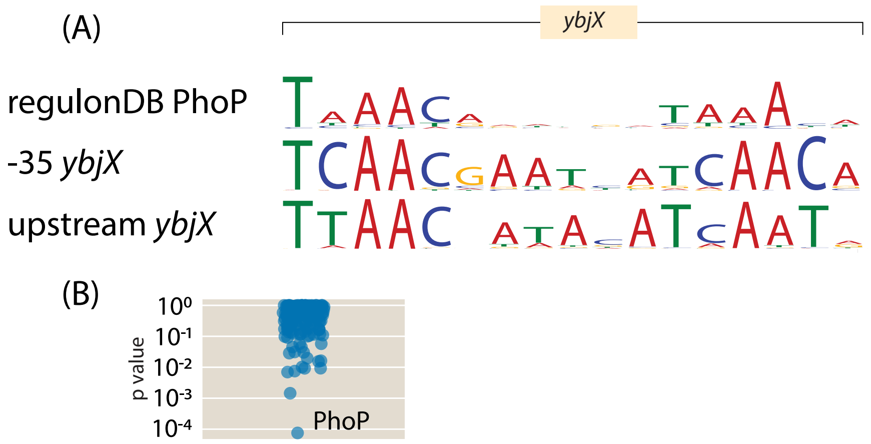

Figure S4: Motif comparison using TOMTOM. Searching our energy motifs against the RegulonDB database using TOMTOM allowed us to guide our TF knockout experiments. Here we show the sequence logos of the PhoP transcription factor from RegulonDB (top) and the one generated from the ybjX promoter energy matrix. E-value = 0.01 using Euclidean distance as a similarity matrix.

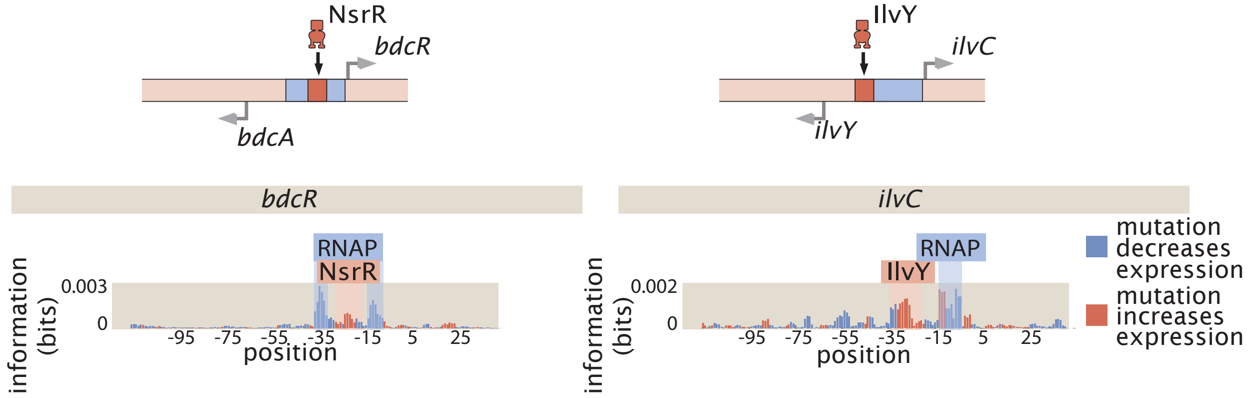

Figure S5: Two cases in which we see transcription factor binding sites that we have found to regulate both of the two divergently transcribed genes.

it also represses the promoter in the absence of that inducer. The production of *ilvC* is known to increase by approximately a factor of 100 in the presence of inducer [16]. The magnitude of the change is attributed to the cooperative binding of two IlvY binding sites, but the lowered expression of the promoter due to IlvY repression in the absence of inducer is also a factor.

#### 4.2 Regulatory cartoons

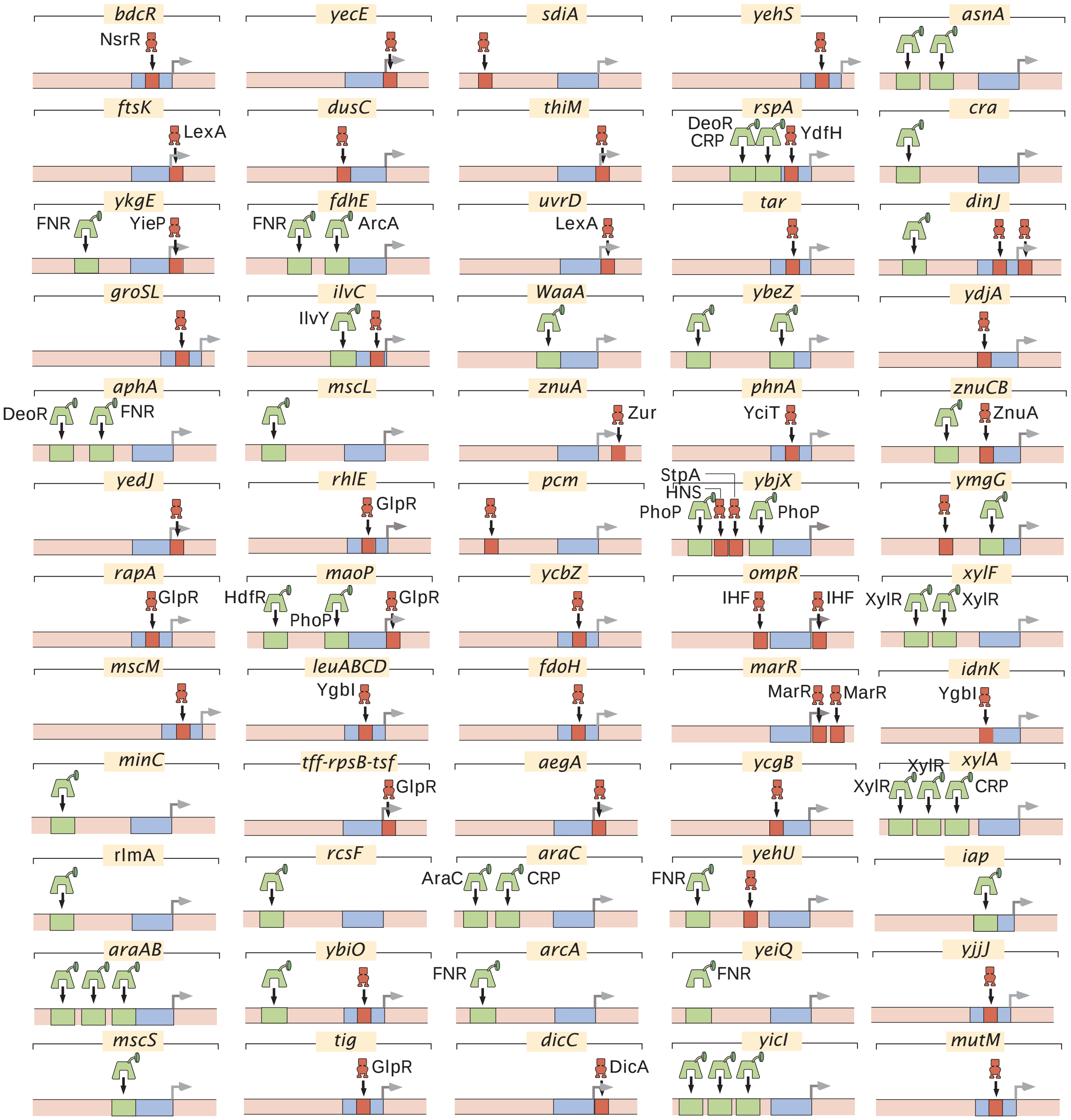

Figure S6: All regulatory cartoons for genes considered in our study.

#### 4.3 Comparison of results to regulonDB

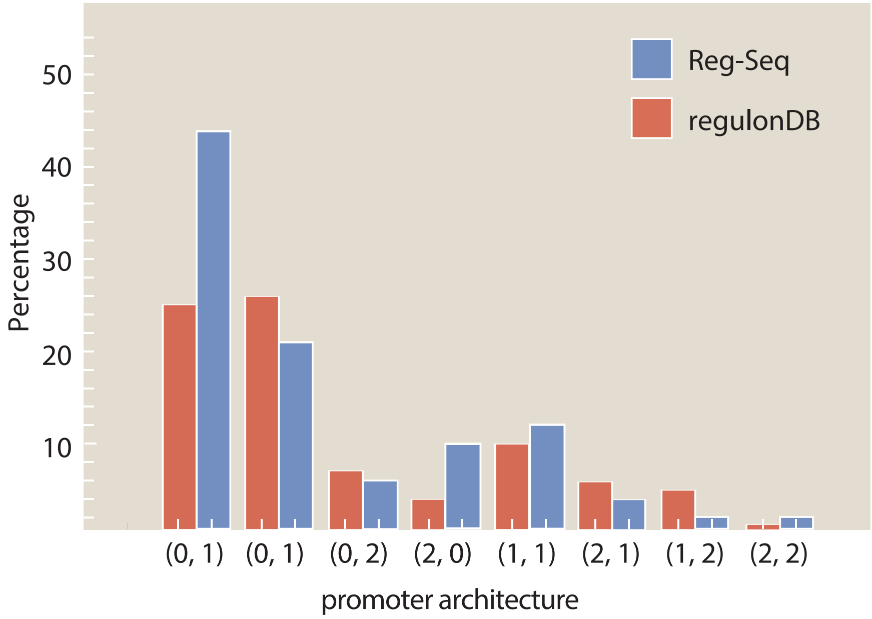

Figure S7: A comparison of the types of architectures found in RegulonDB [6] to the architectures with newly discovered binding sites found in the Reg-Seq study.

[5] Nathan M. Belliveau, Stephanie L. Barnes, William T. Ireland, Daniel L. Jones, Michael J. Sweredoski, Annie Moradian, Sonja Hess, Justin B. Kinney, and Rob Phillips. Systematic approach for dissecting the molecular mechanisms of transcriptional regulation in bacteria.

*Proceedings of the National Academy of Sciences of the United States of America*, 115(21):E4796–E4805, 2018.
